# Concurrent measurement of O_2_ and isoprene production during photosynthesis: pros, cons, and metabolic implications

**DOI:** 10.1101/2023.05.15.540825

**Authors:** Suman Som, Luiza Gallo, Aatish Sunder, Jilian Demus, Tomas F. Dominges, Christina M. Wistrom, Lianhong Gu, Guillaume Tcherkez, Kolby J. Jardine

## Abstract

During oxygenic photosynthesis, oxygen (O_2_) is generated from water photolysis, which provides reducing power to sustain CO_2_ assimilation. To date, traditional leaf gas-exchange experiments have been focused on net CO_2_ exchange (A_net_), with limited observations of net oxygen production (NOP). Here, we present the first gas-exchange/fluorescence system, coupling CO_2_/H_2_O analysis (photosynthesis and transpiration) with NOP and isoprene emission measurements. This configuration allowed us to calculate the assimilatory quotient (AQ = A_net_/NOP) and thus obtain a more complete picture of the photosynthetic redox budget via photosynthetic production of O_2_, electron transport rate (ETR), and isoprene biosynthesis. We used cottonwood leaves (*Populus trichocarpa*) and carried out response curves to light, CO_2_ and temperature along with ^18^O-labelling with ^18^O-enriched water. We found that A_net_ and NOP were linearly correlated across environmental variables with AQ of 1.27 +/- 0.12 regardless of light, CO_2_, and temperature. A_net_ and NOP had optimal temperatures (T_opt_) of 31°C, while ETR (35°C) and isoprene emissions (39°C) had distinctly higher T_opt_. Leaves labelled with H_2_^18^O produced labeled (^18^O^16^O) oxygen with the same T_opt_ as ETR (35°C). The results confirm a tight connection between water oxidation and ETR and are consistent with a suppression of A_net_ and NOP at high temperature driven by an acceleration of (photo)respiration. The findings support the view of isoprene biosynthesis primarily driven by excess photosynthetic ATP/NADPH not consumed by the Calvin cycle during photorespiratory conditions as an important thermotolerance mechanism linked with high rates of CO_2_ and O_2_ recycling.

**Keywords:** Photosynthesis, net oxygen production, gross oxygen production, H_2_^18^O labeling

**One sentence summary:** A leaf gas-exchange system is presented enabling a more complete picture of the photosynthetic redox budget and calculation of the assimilatory quotient.

## Introduction

Terrestrial ecosystems cycle large amounts of carbon dioxide (CO_2_) and oxygen (O_2_) between biosphere and atmosphere via photosynthesis, photorespiration, and respiration. Therefore, plant metabolism represents a critical biological component of the global climate system. For example, when a high resolution (0.05°) dataset for gross primary production (GPP) of sunlit and shaded vegetation canopies from 1992 to 2020 was evaluated, mean global GPP was estimated to be 125.0 ± 3.8 Pg C a^−1^ (Bi et al., 2022). While high rates of photosynthetic CO_2_ assimilation and O_2_ production associated with GPP occur in numerous ecosystems around the world, photorespiration (Wingler et al., 2000) and mitochondrial respiration (Tcherkez et al., 2017), can return a large fraction of assimilated carbon to the atmosphere as CO_2_ while consuming O_2_. For example, terrestrial autotrophic respiration estimates (45–55 Pg C a^−1^) (Luyssaert et al., 2007) represent a major fraction of GPP and are 4.5–6.2 times the average annual CO_2_ release from anthropogenic fossil fuel combustion (8.9–9.9 Pg C/year of CO_2_) over 2008–2017 (Le Quéré et al., 2018).

The majority of gas-exchange observations of photosynthesis and (photo)respiration from individual leaves under controlled environmental conditions have focused on biological and environmental variables impacting net CO_2_ assimilation (A_net_) without the inclusion of net oxygen production (NOP) measurements. This lack of leaf-atmosphere O_2_ flux data is largely due to technical difficulty to measure a small change in O_2_ mole fraction (e.g. 2-200 ppm O_2_) in a high atmospheric O_2_ background (21%, i.e., 210,000 ppm), that is, the high measurement precision needed to clearly resolve relatively small atmospheric O_2_ concentration changes in gas exchange systems (Kim-Hak et al., 2018). Thus, dynamic leaf gas-exchange observations of both A_net_ and NOP as a function of environmental conditions remain rare across diverse plant functional types and ecosystems globally, representing a major knowledge gap in terrestrial ecosystem carbon and oxygen cycling. In addition, most gas-exchange experiments focus on light and CO_2_ response curves in order to obtain photosynthetic parameters important for modeling terrestrial photosynthesis, such as maximum carboxylation velocity catalyzed by Rubisco (V_cmax_) (Sharkey et al., 2007). However, surface temperatures, which are rising at unprecedented rates linked to climate change, have a large impact on leaf CO_2_ and O_2_ metabolism (Sage and Kubien, 2007). Although gross fluxes of photosynthesis (CO_2_ sink, O_2_ source) and (photo)respiration (CO_2_ sources and O_2_ sinks) are known to have different temperature sensitivities (Tcherkez et al., 2017; Dusenge et al., 2019), they remain poorly characterized.

Early studies demonstrated the potential of mass spectrometry to quantify leaf gross production of ^16^O_2_ in the light simultaneously with gross ^18^O_2_ uptake under a recirculated leaf headspace atmosphere of 21% ^18^O_2_ (Canvin et al., 1980). This technique was used with potted tomato plants to demonstrate that during leaf water stress, gross oxygen uptake and consumption decline, suggesting that photosystem II, the Calvin cycle, and mitochondrial respiration are down-regulated (Haupt-Herting and Fock, 2002). More recently, a new method based on measuring δ^18^O of O_2_ in air of a detached leaf pre-treated with H_2_^18^O was used to estimate gross oxygen production (GOP) and NOP determined separately from the increase in the O_2_/N_2_ ratio (Gauthier et al., 2018). Thus, this method has the potential to directly determine the optimal temperature (T_opt_) of GOP, which is expected to follow the pattern of photosynthetic electron transport rates (ETR) determined from chlorophyll fluorescence.

While relatively few observations have been reported, the interface of environmentally controlled dynamic leaf chambers to high precision real-time oxygen sensors has opened the door to concurrent measurement of A_net_, net oxygen production (NOP), and thus the apparent assimilatory CO_2_/O_2_ quotient (AQ = A_net_/NOP) (Cousins and Bloom, 2003). AQ is expected to be near 1.0 when the Calvin cycle is the dominant sink of photosynthetic energy and reducing equivalents (and when carbohydrates are used as the respiratory substrate). However, AQ can deviate from 1.0 linked to the activity of alternate sinks not directly coupled to CO_2_/O_2_ metabolism including nitrate photo-assimilation (Smart and Bloom, 2001), and to a lesser extent, lipid and lignin biosynthesis (Cen et al., 2001; Cousins and Bloom, 2003; Searles and Bloom, 2003). Isoprene is a volatile light-dependent photosynthetic lipid product produced and emitted by leaves of many tree species globally (Jardine et al., 2020). Isoprene is not stored in plants, with leaf emissions directly linked to production rates within chloroplasts via the isoprenoid pathway through the utilization of both products from the ‘light’ reactions such as reducing power (NADPH) and energy (ATP), and triose phosphate carbon skeletons from the ‘dark’ reactions (Rodrigues et al., 2020). While the majority of carbon in isoprene derive from atmospheric CO_2_ within minutes of photosynthesis in the light (Karl et al., 2002), alternate non-photosynthetic carbon sources for isoprene increase during stress (Funk et al., 2004) such as high temperature (Jardine et al., 2014). Externally supplied pyruvate has been demonstrated as an effective isoprene carbon source (Jardine et al., 2010) and studies that labeled leaf isoprene with ^13^CO_2_ suggested that pyruvate for isoprene synthesis may derive from both recent photosynthesis and cytosolic stored carbon sources like glycolysis (Karl et al., 2002). However, recent studies using CO_2_-free air suggested that internal CO_2_ and O_2_ recycling under photorespiratory conditions may play important roles as an ‘alternative’ carbon source for isoprene (Garcia et al., 2019). As internal CO_2_ and O_2_ recycling in leaves do not alter net leaf-atmosphere fluxes of O_2_ and CO_2_, they are difficult to study (Tcherkez et al., 2017), but known to accelerate under stress (Ma et al., 2001). Thus, isoprene emissions may provide insight into the role of internal CO_2_ and O_2_ recycling in leaves under photorespiratory stress conditions such as drought, high light, and high temperature (Voss et al., 2013).

Here, we coupled a high precision O_2_ cavity ring down spectrometer (CRDS) and a proton transfer reaction-mass spectrometer (PTR-MS) to the sub-sampling port of a commercial gas exchange system with full environmental control and integrated fluorimeter (Li-6800, with the 6-cm^2^ leaf chamber). To our knowledge, this coupling represents the first CO_2_ gas exchange (with chlorophyll fluorescence)/O_2_ measurements/isoprene analysis system for simultaneous, real-time quantification of photosynthetic parameters such as electron transport rate (ETR), net CO_2_ assimilation (A_net_), transpiration, net oxygen evolution (NOP), and isoprene emission. Furthermore, it allows the measurements of the assimilatory quotient (AQ = A_net_/NOP) so as to get information on the photosynthetic redox budget in leaves. We carried out response curves to photosynthetically active radiation (PAR), leaf internal CO_2_ concentrations (C_i_), and leaf temperature, using mature leaves of black cottonwood (*Populus trichocarpa* Torr. & Gray).

We hypothesized that gross fluxes of photosynthesis, (photo)respiration, and isoprenoid synthesis have distinctly different temperature sensitivities and optimum temperatures. This would imply that as leaf temperature increases, an increasing proportion of ATP and NADPH from ‘light’ reactions are used for photorespiration (Long, 1991) and isoprenoid synthesis (Rodrigues et al., 2020) instead of CO_2_ assimilation. Due to partial stomatal closure leading to reduced C_i_, the suppression of atmospheric CO_2_ uptake is partially compensated for by increased refixation of (photo)respiratory CO_2_ at high temperature (Voss et al., 2013) and thus enhanced CO_2_/O_2_ recycling (Garcia et al., 2019). In addition, we can expect high isoprene emission due to excess NADPH and ATP not being used by the Calvin cycle (Morfopoulos et al., 2014) and the relatively high temperature optimum of isoprene synthase (e.g. 40-45 ℃, depending on growth temperature) (Monson et al., 1992). To test this hypothesis, we switched the CRDS to the isotope mode which enables direct measurement of the δ^18^O value in O_2_, with a precision better than 1‰ using 5-min averages. By supplying detached poplar leaves a solution of ^18^O-enriched water via the transpiration stream (cut petiole on the outside of the leaf chamber instead of inside the chamber, i.e. we modified the method of Gauthier *et al*. (2018), with minimal leaks around the leaf/petiole using a commercial silicon-based gasket integrated into a large leaf chamber, 36 cm^2^ under controlled actinic lighting and leaf temperature. The large leaf chamber was found to be important to detect the ^18^O-enrichment in O_2_ evolved by photosynthesis, since it allows increased gas residence time in the chamber and higher O_2_ inlet-outlet gradient.

## Material and Methods

### Plant material

We used 15 potted California poplar (*Populus trichocarpa*) saplings (average height of 2 m) obtained from Plants of the Wild (Washington State, USA) and maintained for three years in the South Greenhouse at the Oxford Tract Experimental Facility in Berkeley, CA, USA, where they were regularly watered and subject to standard pest control practices. Water was delivered to each individual using an automated watering system in #15 pots containing Supersoil planting media (Scotts Co., Marysville, Ohio, USA). Nitrogen in the form of both nitrate (NO_3_^-^) and ammonium (NH_4_^+^) was supplied using three fertilizers including slow release Osmocote plus (15-9-12) added directly to the soil during potting (240 g per #15 pot) and Yara Liva CaNO_3_ at 90 ppm and Peters Professional 20-20-20 at 74 ppm, mixed together in the irrigation water, applied 5 times per week to soil saturation. Ambient natural light was supplemented with LED lighting using an Argus Titan environmental control system with photocell (Argus Controls, British Columbia, Canada). The controller was programmed to turn LED lights off when detecting exterior light levels above 850 μmol m^-2^ s^-1^ during the 16-hour photoperiod (6:00 AM to 10:00 PM). Poplar branches were detached from one of the 15 trees in the greenhouse in the morning (9:00-12:00), with stems immediately immersed and recut under water, and then transferred to the nearby laboratory. The selected leaf to be studied with gas exchange was placed in the leaf chamber, ensuring complete coverage of the 6-cm^2^ or 36-cm^2^ surface area, depending on the leaf chamber used. To hydrate the branch and minimize water loss through transpiration, the branch outside the leaf chamber was immediately covered with a mylar sheet with wet paper towels placed around the base. Therefore, only the leaf in the chamber was actively transpiring. This was found to be important at high leaf temperatures (e.g., 40 ℃) to avoid leaf desiccation in the chamber associated with elevated transpiration rates. After an acclimation period (15 min), leaf response curves to PAR, C_i_, or leaf temperature were initiated (**Figure 1a**). In a separate set of experiments with only the large leaf chamber, following the installation of a leaf in the chamber in darkness, the petiole was cut and placed in a solution of H_2_^18^O for a pre-treatment period before a leaf temperature response curve was initiated (**Figure 1b**).

**Figure 1.**
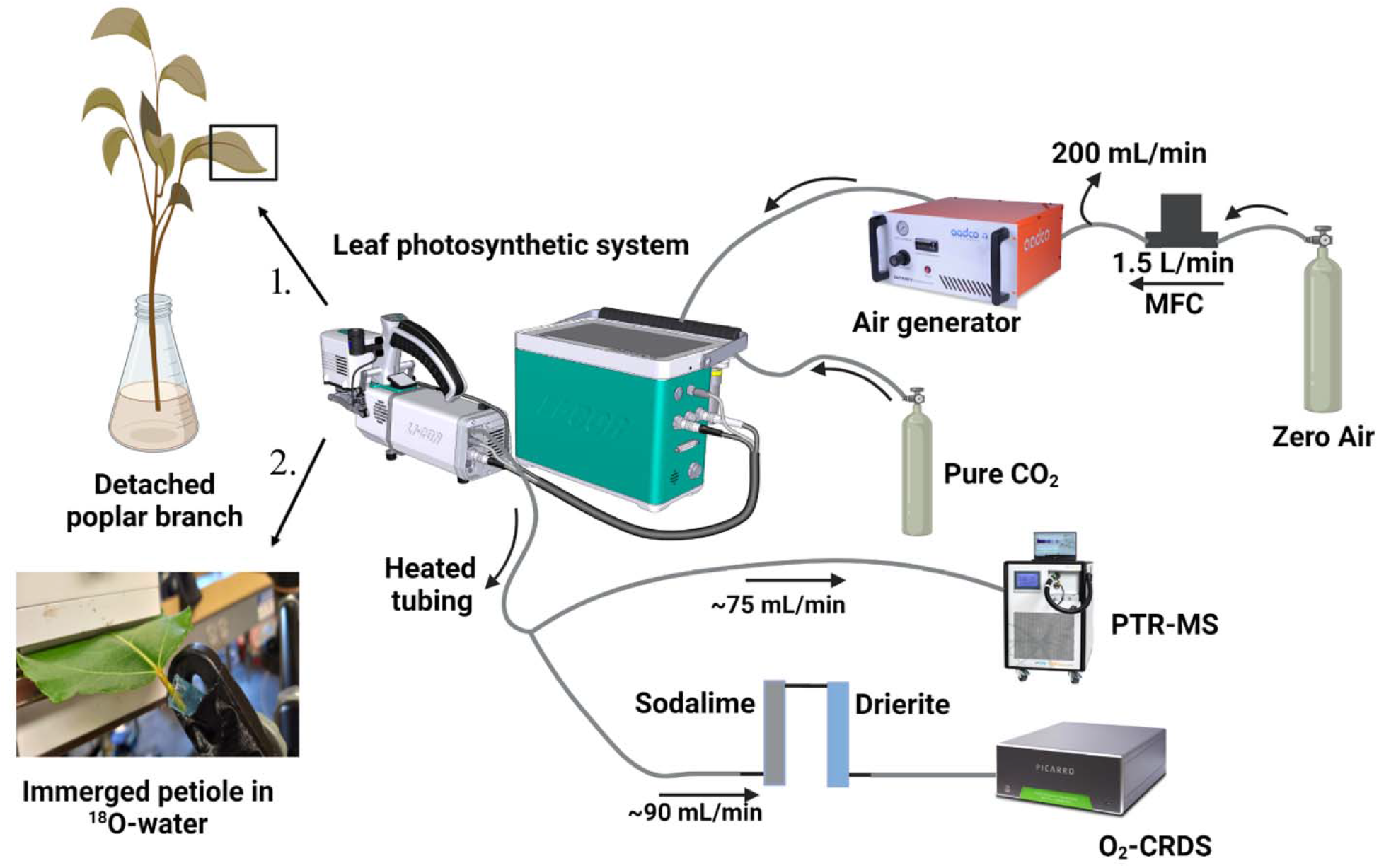
Schematic diagram of experimental setup for (**1**) real-time leaf to atmosphere fluxes of CO_2_, H_2_O, O_2_, and isoprene together with *chlorophyll fluorescence across environmental leaf response curves of PAR, C_i_, and leaf temperature using both small and large leaf chambers (**2**) real-time leaf to atmosphere fluxes of CO_2_, H_2_O, and isoprene together with *δ^18^O of leaf chamber O_2_ during leaf temperature response curves using the large leaf chamber. *chlorophyll fluorescence was only quantified using the small leaf chamber.

### Leaf gas exchange system

For all experiments, gas exchange (CO_2_ and H_2_O) was carried out under controlled environmental conditions using a portable photosynthesis system (Li-6800, Li-COR Biosciences, Nebraska, USA) coupled to a high precision O_2_ CRDS (Picarro Inc. California, USA) and quadrupole PTR-MS (Ionicon, Austria) for headspace O_2_ and isoprene concentration measurements, respectively (**Figure 1**). A fraction of air exiting the leaf chamber was diverted from the Li-6800 subsampling gas port to the O_2_ CRDS (90 mL min^-1^) and PTR-MS (75 mL min^-1^) using a ⅛” O.D. Teflon PTFE tube that was continuously heated to 50-60 ℃ with self-regulating heating tape (SLR10, Omega Engineering, USA) to prevent condensation and gas-tubing wall interactions prior to gas analysis by the CRDS and PTR-MS sensors. The gas source for the Li-6800 was supplied externally by overblowing a tee fitting with high purity zero air (ultra-zero air, CAS: 132259-10-0, Linde Gas) such that at least 200 mL min^-1^ vented externally while the remaining flow (1,300 mL min^-1^) was passed through a platinum catalytic converter held at 280 ℃ (ZA30 catalyst, Aadco instruments, USA) to oxidize any trace VOCs before entering the air inlet of the Li6800. Therefore, inlet air delivered to the leaf chamber was CO_2_-, H_2_O-, and VOC-free while maintaining a constant concentration of O_2_, which varied from cylinder to cylinder between 20.09 and 21.03%. Leaf chamber humidity was regulated by balancing air flow through the desiccant (Drierite with 10-20 mesh size CAS:778-18-9, Drierite) and humidifier (⅛” O.D. Nafion tubing immersed in ACS/HPLC water, CAS: 7732-18-5, Honeywell) in order to maintain the absolute humidity of the reference air at the desired setpoint. CO_2_ mole fraction inside chamber was controlled by passing all airflow through the CO_2_ scrubber (indicating soda lime, 4-8 mesh size, CAS: 8006-28-8, Thermo Scientific) while carbon dioxide was supplied by an external cylinder (CAS 124-38-9, 99.9% CO_2_, Praxair). When LED light inside the leaf chamber was switched on, the spectrum was set to r90b10 (990 µmol m^-2^ s^-1^ red, 10 µmol m^-2^ s^-1^ blue).

Leaf isoprene emission was measured for all gas-exchange measurements using a real-time high sensitivity quadrupole proton transfer reaction mass spectrometry (PTR-MS, with a QMZ 422 quadrupole, Balzers, Switzerland) as previously described (Jardine et al., 2014). The PTR-MS was operated with a drift tube voltage of 440 V and pressure of 1.8 mbar. For each measurement cycle lasting 24 sec, the following mass to charge (m/z) ratios were monitored: m/z 21 (H_3_^18^O^+^), m/z 37 (H_3_O^+^-H_2_O), and m/z 69 (protonated isoprene: H^+^-C_5_H_8_).

Two different leaf chambers (small and large) were used with distinct advantages and disadvantages (**Figure 2**). The small leaf chamber (6 cm^2^) had the added advantage of including an integrated chlorophyll fluorometer (6800-01A, LI-COR Biosciences, USA) together with H_2_O and CO_2_ gas exchange. Due to the rerouting of a fraction of the outlet air for simultaneous measurements of O_2_ and isoprene concentrations, and small leaks that formed between the gasket and the leaf/petiole, an optimized flow rate of 323-363 mL min^-1^ (240-270 µmol mol^-1^) was delivered to the leaf chamber which ensured high O_2_ gradients while maintaining sufficient flow for O_2_ (90 mL min^-1^), isoprene (75 mL min^-1^), and CO_2_ + H_2_O (158-198 mL min^-1^) measurements (**Figure 2a**). Chlorophyll fluorescence data were simultaneously recorded for each of the three environmental variables gas-exchange experiments (light, CO_2_ and temperature responses curves) using an integrated multiphase flash fluorometer system (model 6800-01A, LI-COR Biosciences, USA) using an actinic light pulse of 1,000 μmol m^-^² s^-^¹ PAR. The light pulse was applied for 1,000 ms with a dark modulation rate of 50 Hz, light modulation rate of 1 kHz, flash modulation rate of 250 kHz, with 15 seconds chlorophyll fluorescence signal averaging (100 Hz data output rate with a margin of 5 points before and after flash). Photosynthetic electron transport rate (ETR, µmol e^-^ m^-2^ s^-1^) was calculated.

**Figure 2:**
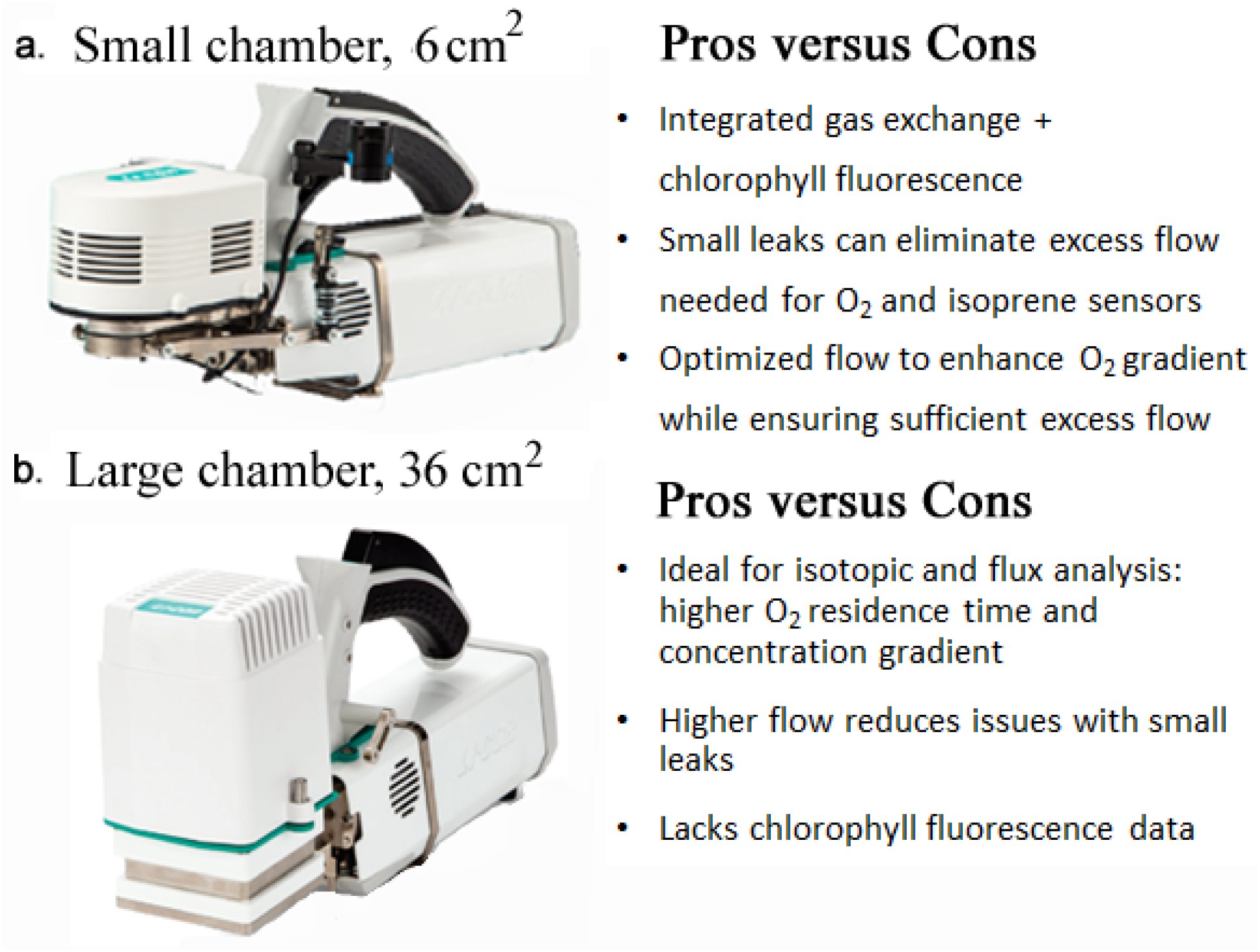
Pros and Cons of quantifying leaf net O_2_ production (NOP) fluxes and δ^18^O of leaf headspace O_2_ using **a**) a small dynamic leaf chamber (6 cm^2^) with integrated chlorophyll fluorimeter and **b**) a large dynamic leaf chamber with actinic light source (36 cm^2^).

A large leaf chamber with integrated LED light source (model 6800-03 LI-COR Biosciences, USA) was also used with the advantage of enclosing a much larger area of enclosed leaf (36 cm^2^). This allowed for a higher flow rate of air to be delivered to the leaf chamber (538 mL min^-1^ or 400 µmol s^-1^). While lacking a fluorimeter, the large leaf chamber can control leaf temperature, the actinic light spectra (r90b10), and PAR intensity identically to the small chamber (**Figure 2b**).

### Photosynthetic and ETR response curves

#### Light response curves

Photosynthetic light response curves were carried out at constant leaf temperature (32 °C), leaf chamber CO_2_ mole fraction of 400 µmol mol^-1^, and reference (inlet) air humidity of 12 mmol mol^-1^. For both large and small chambers, after 30 min dark acclimation (PAR: 0 µmol m^-2^ s^-1^), leaf gas exchange and chlorophyll fluorescence (small chamber only) responses to light intensity was initiated. This included a sequence of increasing followed by decreasing PAR (0, 200, 400, 600, 800, 1000, 1200, 1400, 1600, 1200, 800, 400, 100, 50, 40, 30, 20, 10 and 0 µmol m^-2^ s^-1^). Total time duration for the light response curves was 200 min. Three replicate light response curves were collected using the small chamber with gas exchange and chlorophyll fluorescence and 2 replicate light response curves were collected using the large chamber with gas exchange only. For each replicate, a branch from a different tree (5 out of 15 total) was used.

#### C_i_ response curves

The response of leaf gas exchange to intercellular CO_2_ mole fraction (C_i_) (i.e., A_net_/C_i_ response curves) were carried out by varying leaf chamber CO_2_ mole fraction (C_a_) while maintaining constant leaf temperature (32 °C), PAR (1,000 µmol m^-2^ s^-1^) and reference (inlet) air humidity (10 mmol mol^-1^). For both large and small leaf chambers, after 30 min light acclimation (PAR: 1,000 µmol m^-2^s^-1^), leaf gas exchange and chlorophyll fluorescence (small chamber only) response curves to CO_2_ were initiated. This included a sequence of decreasing followed by increasing CO_2_ (400, 350, 300, 250, 200, 150, 125, 100, 75, 50, 25, 0, 50, 100, 150, 200, 250, 300, 400, 500, 600, 700, 800, 900, 1000 ppm). Total time duration for the C_i_ response curves was 160 min. Four replicate C_i_ response curves were collected using the small chamber and 3 replicate C_i_ response curves were collected using the large chamber. For each replicate, a branch from a different tree (7 out of 15 total) was used.

#### Leaf temperature response curves

Leaf temperature response curves were carried out with varying leaf temperature at constant C_a_ (400 µmol mol^-1^) and inlet (reference) air humidity (0-8 mmol mol^-1^). To prevent condensation in the large leaf chamber, dry inlet air (with 0 mmol mol^-1^ water vapor) was supplied, while in the small leaf chamber, the shorter gas residence time allowed us to use inlet air with a humidity of 8 mmol mol^-1^ (see also *Discussion*). For both large and small leaf chambers, after 30 min dark acclimation (PAR: 0 µmol m^-2^ s^-1^) at 25 °C leaf temperature, leaf gas exchange (both chambers) and chlorophyll fluorescence responses (small chamber only) to leaf temperature commenced. The sequence started with leaf dark respiration measurements at 25.0 °C (PAR: 0 µmol m^-2^ s^-1^). Following a period of light acclimation, the temperature response curve in the light (PAR: 1,000 µmol m^-2^ s^-1^) was initiated with increasing leaf temperatures (25.0, 27.5, 30.0, 32.5, 35.0, 37.5, 40.0 °C). After the temperature response curve, incident light was switched off to record leaf dark respiration at 40°C. Total time duration for leaf temperature response curves was 120 min. Eight replicate leaf temperature response curves were collected using the small chamber. For each replicate, a branch from a different tree (8 out of 15 total) was utilized. In addition, seven replicate leaf temperature response curves were collected using the large chamber (1 branch from 7 trees).

### Leaf H_2_^18^O labeling

To determine optimal temperature of gross oxygen production (GOP), leaf responses to temperature were monitored using the large leaf chamber with detached poplar leaves pre-treated with a solution of H ^18^O water (7 replicate temperature curves, 1 leaf per tree). Water enriched in H_2_^18^O (δ^18^O value of +8,000‰ vs SMOW) was prepared by diluting a 10 atom % H_2_^18^O (CAS:14314-42-2, Sigma-Aldrich) with HPLC grade water. The leaf was detached from the branch and the petiole immediately recut under H_2_^18^O enriched water, and then placed in the large chamber under constant light (PAR: 1,000 µmol m^-2^s^-1^), leaf temperature (32.0 °C), and CO_2_ mole fraction (400 µmol mol^-1^). δ^18^O of O_2_ (outlet air) was measured before, during, and after gas exchange experiments with a leaf. Concurrently, continuous measurement of leaf isoprene emission and the ^18^O/^16^O ratio of transpired water vapor were carried out using PTR-MS (**Figure 1b**). Pre-treatment of the leaf with the H ^18^O enriched water occurred for 2-3 h during which the δ^18^O values reached a steady state, indicating the turn-over of all non-static leaf water pools. Following the pre-treatment period, the leaf temperature curve was carried out with the same protocol as that used for intact leaves (see above).

### Real-time measurement of leaf net oxygen production (NOP) and δ^18^O

The infrared laser-based cavity ring-down spectrometer (CRDS, Picarro G2207, O_2_/H_2_O, Santa Clara, USA) was used for continuous high precision measurement of oxygen mole fraction and δ^18^O values in O_2_. In fact, the CRDS spectrometer could be operated in two different modes: high precision concentration and isotopic ratio modes. In concentration mode, O_2_ mole fraction was measured with < 2 ppm precision using 7-min averages. O_2_ reference mole fraction measurements were made with an empty chamber before and after all leaf environmental response curves and used to calculate the change in O_2_ concen trations due to leaf gas exchange (ΔO_2_). Small drifts in O_2_ inlet mole fraction during the response curve were < 20 ppm O_2_. This drift in reference leaf chamber O_2_ concentrations is attributed to the CRDS itself and was subtracted from the headspace O_2_ concentrations when a leaf was in the chamber. That way, the difference in O_2_ mole fractions between leaf chamber and reference air (ΔO_2_) could be determined in real-time during the environmental response curves. Leaf NOP (µmol m^-2^ s^-1^) fluxes were calculated using **Eq. 1** where µ is the air flow rate entering the leaf chamber (mol air s^-1^), ΔO_2_ is the difference in oxygen mole fraction between leaf chamber and reference air corrected for CRDS drift (µmol mol^-1^), and S is leaf surface area (0.0006 and 0.0036 m^2^) placed inside the chamber. Note, that due to quantitative scrubbing of H_2_O and CO_2_ from outlet air just prior to making O_2_ measurements by the CRDS, corrections associated with air flow rate due to transpiration and photosynthesis were not necessary.

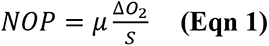

To determine the average AQ for each environmental leaf response curve, a linear regression analysis was performed with A_net_ (y-axis) plotted against NOP (x-axis). Highly linear correlations were observed in all cases, with the slope of the regression representing AQ = A_net_/NOP. In O_2_ CRDS isotope mode, δ^18^O of outlet air (without or with a leaf) was measured with <1‰ precision using 7-min averages.

## Results

### Light response

Light response curves are shown in **Figure 3** and supplementary **Figures S1-S10**. A_net_ showed a slightly higher magnitude in both the dark (more negative) and in the light (more positive) relative to NOP (**Figure 3b)**. This resulted in an assimilatory quotient AQ (= A_net_/NOP) slightly higher than unity (AQ = 1.3 in **Figure 3c**). For example, dark CO_2_ evolution was 3.8 µmol m^-2^ s^-1^ while dark oxygen consumption was 2.5 µmol m^-2^ s^-1^. Similarly, under saturating light (1,600 µmol m^-2^s^-1^ PAR), A_net_ (18.4 µmol m^-2^s^-1^) was slightly higher than NOP (15.5 µmol m^-2^ s^-1^) (**Figure 3b,c)**. Both A_net_ and NOP increased with PAR. At low light intensity (0-200 µmol m^-2^ s^-1^) a linear response was observed with A_net_, NOP, and ETR (**Figure 3b** and supplementary **Figures S1-S10)**. Stomatal conductance (g_s_) and transpiration (E) also increased with PAR, reaching a maximum value (g_s_: 0.35 mol m^-2^s^-1^, E: 8.0 mmol m^-2^s^-1^) at 1,200 µmol m^-2^ s^-1^. At high light, both g_s_ and E decreased slightly. AQ values determined using the small leaf chamber (6 cm^2^, AQ = 1.26 +/- 0.06) were similar to those determined with the large leaf chamber (36 cm^2^, AQ = 1.22 +/- 0.01) (**Table 1**).

**Figure 3.**
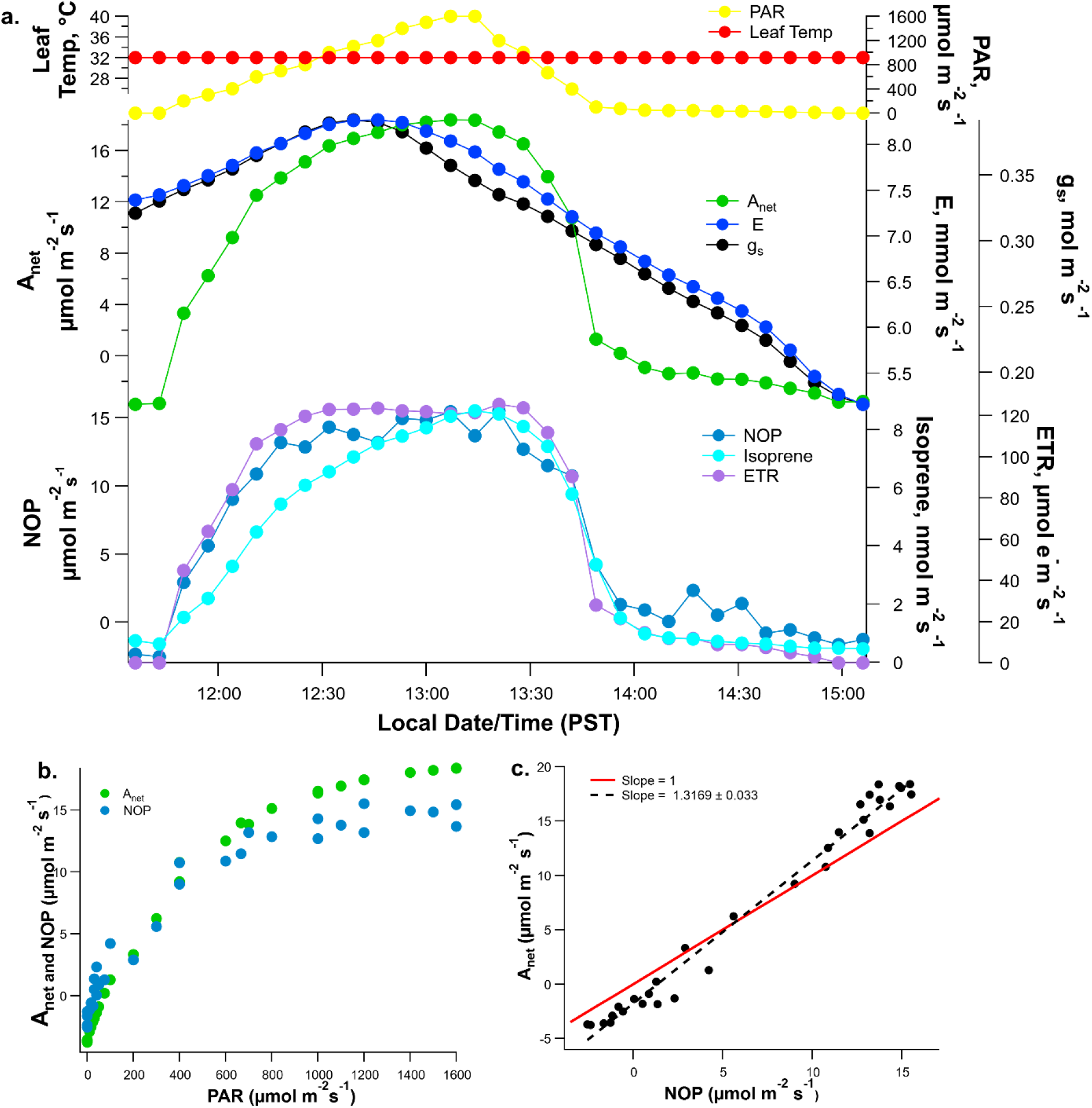
**a.** Example real-time leaf-gas exchange fluxes of A_net_, NOP, E, and isoprene emissions together with chlorophyll fluorescence-derived ETR during controlled light response curves (photosynthetically active radiation, PAR) under constant leaf temperature (32 °C) and leaf chamber headspace CO_2_ concentrations (400 ppm) collected using the 6 cm^2^ leaf chamber with integrated chlorophyll fluorometer. **b.** A_net_ and NOP plotted as a function of PAR. **c.** Linear regression between A_net_ and NOP. Note the slope of the regression as well as the 1:1 line.

**Table 1:** Assimilatory quotients (AQ = A_net_/NOP) determined as the slope from linear regressions between A_net_ and NOP during leaf gas-exchange response curves to light (PAR), leaf internal CO_2_ concentrations (C_i_) and leaf temperature for both large and small dynamic leaf chambers. AQ values shown for are mean ± 1 standard deviation with N indicating the number of replicate leaf response curves.

| Leaf Chamber | AQ (Light) | AQ (C<sub>i</sub>) | AQ (Temp) | AQ (All) |
| --- | --- | --- | --- | --- |
| Large chamber | 1.22 ± 0.01 (N= 2) | 1.27 ± 0.02 (N= 3) | 1.40 ± 0.16 (N= 7) | 1.34 ± 0.14 (N= 12) |
| Small chamber | 1.26 ± 0.06 (N= 3) | 1.23 ± 0.08 (N= 4) | 1.21 ± 0.09 (N= 8) | 1.23 ± 0.08 (N= 15) |
| Large + Small chamber | 1.25 ± 0.04 (N= 5) | 1.25 ± 0.06 (N= 7) | 1.30 ± 0.15 (N= 15) | 1.27 ± 0.12 (N = 27) |

### CO_2_ response

Response curves to intercellular CO_2_ mole fraction (C_i_) are shown in **Figure 4** and supplementary **Figures S11-S24**. Both A_net_ and NOP increased with C_i_, although A_net_ showed a slightly larger magnitude at both low (more negative) and high C_i_ (more positive) than NOP (**Figure 4b)**. AQ determined using the small leaf chamber was slightly higher than unity (AQ = 1.3 for the leaf shown in **Figure 4c**). Across all C_i_ response curves, AQ values determined using the small leaf chamber (6 cm^2^, AQ = 1.23 +/- 0.08) were similar to those determined using the large leaf chamber (36 cm^2^, AQ = 1.27 +/- 0.02) (**Table 1**). As atmospheric CO_2_ declined from 400 to 0 µmol mol^-1^, C_i_ declined from 320 to 31 µmol mol^-1^. A_net_ and NOP became negative below 56 µmol mol^-1^. Below the compensation point, at the lowest C_i_ (31 µmol mol^-1^), CO_2_ evolution (3.8 µmol m^-2^s^-1^) was slightly higher than oxygen consumption (2.2 µmol m^-2^s^-1^). In contrast, isoprene emissions were stimulated as C_i_ declined from 320 to 56 µmol mol^-1^ (+67%), followed by a decline as C_i_ reached the lowest value, 31 µmol mol^-1^ (-19%). A photosynthetic plateau occurred for C_i_ above 680 µmol mol^-1^ while isoprene emissions were suppressed at elevated C_i_ (-87% from 207 to 868 µmol mol^-1^). Taken as a whole, while A_net_, NOP, and ETR showed very strong dependency on C_i_ while changes in isoprene emissions were relatively small.

**Figure 4.**
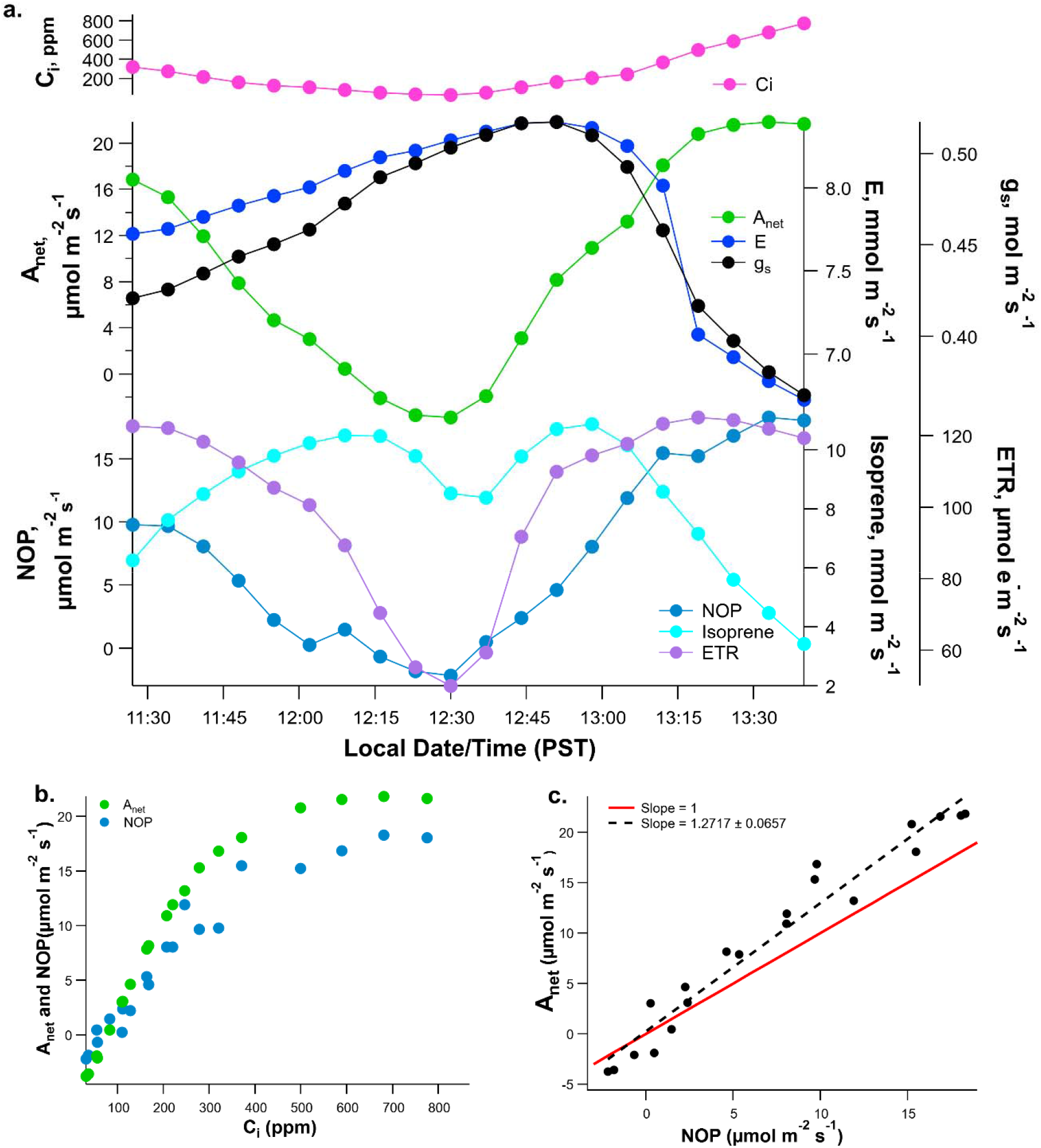
(a). Example real-time leaf-gas exchange fluxes of A_net_, NOP, E, and isoprene emissions together with ETR during controlled C_i_ (photosynthetically active radiation, PAR) response curves under constant leaf temperature (32 °C) and PAR (1000 µmol m^-2^ s^-1^) collected using the 6 cm^2^ leaf chamber with integrated chlorophyll fluorimeter. (b). A_net_ and NOP plotted as a function of C_i_. (c) Linear regression between A_net_ and NOP. Note the slope of the regression as well as the 1:1 line.

### Temperature response

Leaf temperature response curves are shown in **Figure 5** and supplementary **Figures S25-S54**. In darkness, CO_2_ evolution at 25°C (1.8 µmol m^-2^ s^-1^) was higher than oxygen consumption (0.2 µmol m^-2^ s^-1^). Isoprene emissions were negligible in the dark. However, high values of stomatal conductance g_s_ (0.26 mol m^-2^ s^-1^) and transpiration E (3.2 mmol mol^-1^) were observed. In the light at 25 °C, A_net_, NOP, and ETR showed high values while isoprene emissions remained low, but detectable (< 0.5 nmol m^-2^ s^-1^). As leaf temperature increased in the light, A_net_ and NOP reached maximum values near 31 °C and then decreased slightly at higher temperature. In contrast, ETR continued to increase in the light to a maximum near 36 °C while isoprene emissions continued to increase up to the highest leaf temperature used (40 °C). Upon switching off the light at the highest leaf temperature (40 °C), A_net_ and NOP rapidly declined and became negative while isoprene emission was nearly suppressed. In all leaves studied, an increase in leaf dark respiration (A_net_ and NOP) was observed at 40 °C relative to 25 °C. Similar to what was observed with light and C_i_ response curves, leaf temperature response curves resulted in A_net_ value slightly larger than NOP (**Figure 5b),** that is, AQ values slightly higher than unity (AQ = 1.4 in **Figure 5c**). AQ values determined using the small leaf chamber (6 cm^2^, AQ = 1.21 +/- 0.09) were similar to those determined from the large leaf chamber (36 cm^2^, AQ = 1.40 +/- 0.16) (**Table 1**).

**Figure 5.**
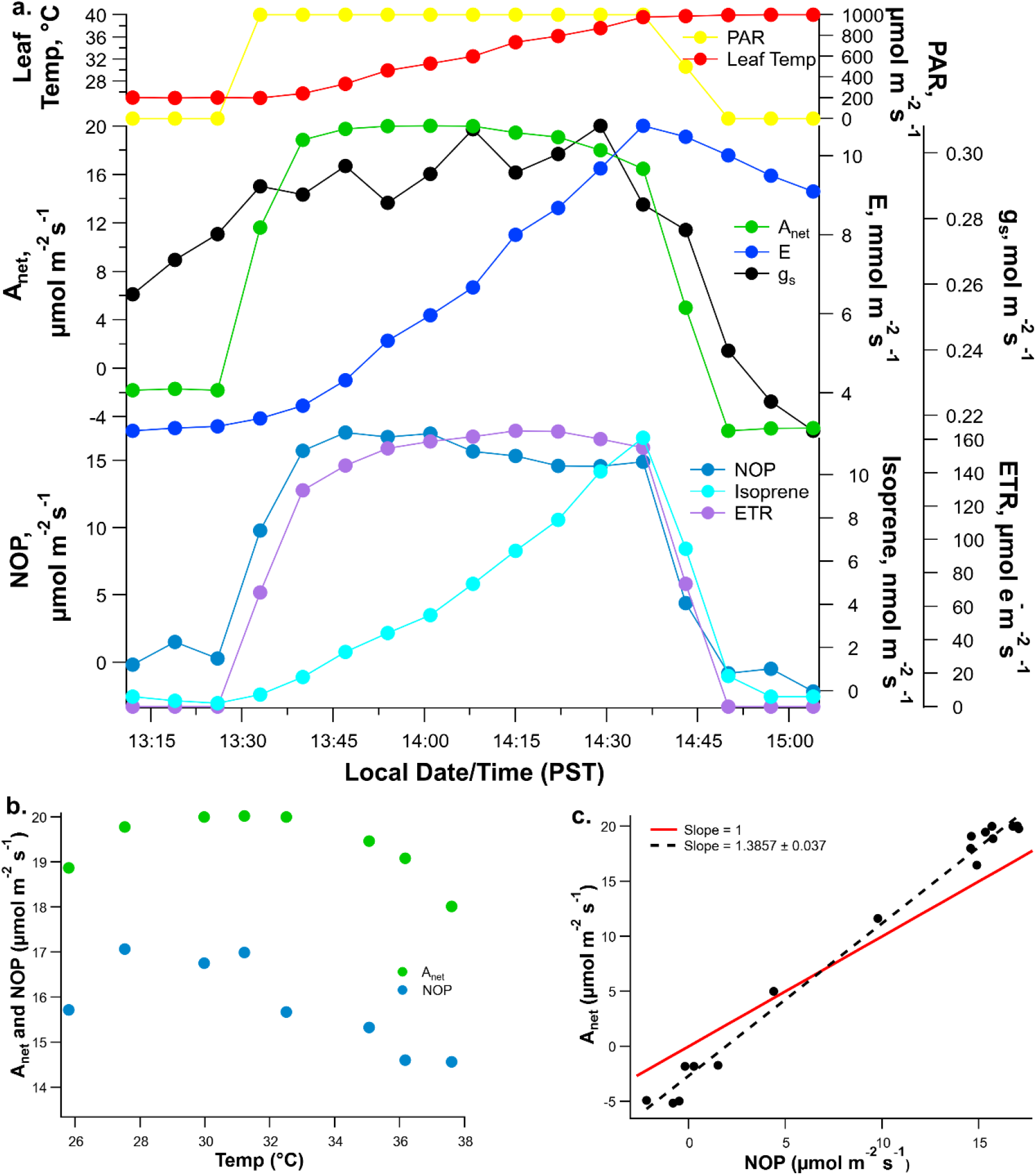
(a). Example, real-time leaf-gas exchange fluxes of A_net_, NOP, E, and isoprene emissions together with ETR during controlled leaf temperature response curves under constant leaf headspace enclosure CO_2_ (400 ppm) and PAR (1000 µmol m^-2^ s^-1^) collected using the 6 cm^2^ leaf chamber with integrated chlorophyll fluorimeter. (b). A_net_ and NOP plotted as a function of leaf temperature. (c) Linear regression between A_net_ and NOP. Note the slope of the regression as well as the 1:1 line shown.

### Temperature response curves with ^18^O-water

The temperature dependence of gross oxygen production (GOP) was assessed using detached mature poplar leaves placed in the large (36 cm^2^) chamber in the light, with the petiole immersed in ^18^O-enriched water (**Figure 6** and supplementary **Figures S55-S68**). During the pre-treatment period (incubation in ^18^O-enriched water) in the light at 30 °C, the δ^18^O of outlet O_2_ increased from the background value (ca. +11‰) and reached +21‰ within two hours in the steady state (**Figure 6a**). During this pre-treatment period, leaf isoprene emissions also increased, but reached a steady state much faster (within 15 min). Upon switching off the light and reducing leaf temperature to 25 °C, δ^18^O of outlet O_2_ values quickly returned to background values of +12‰. When the light was switched on again at 25 °C, δ^18^O values reached +20‰ and increased with leaf temperature up to a maximum of +23‰ at 37.5 °C, and then decreased slightly at the highest leaf temperature (40.0 °C). When the light was switched off at 40 °C (end of the temperature response curve), δ^18^O of outlet O_2_ rapidly returned to the background value of +12‰. Although A_net_ showed a similar optimum leaf temperature of (30 °C) to that of non-detached leaves (**Fig. 5b** versus **6b**), the optimum temperature of δ^18^O was much higher (37.5 °C).

**Figure 6.**
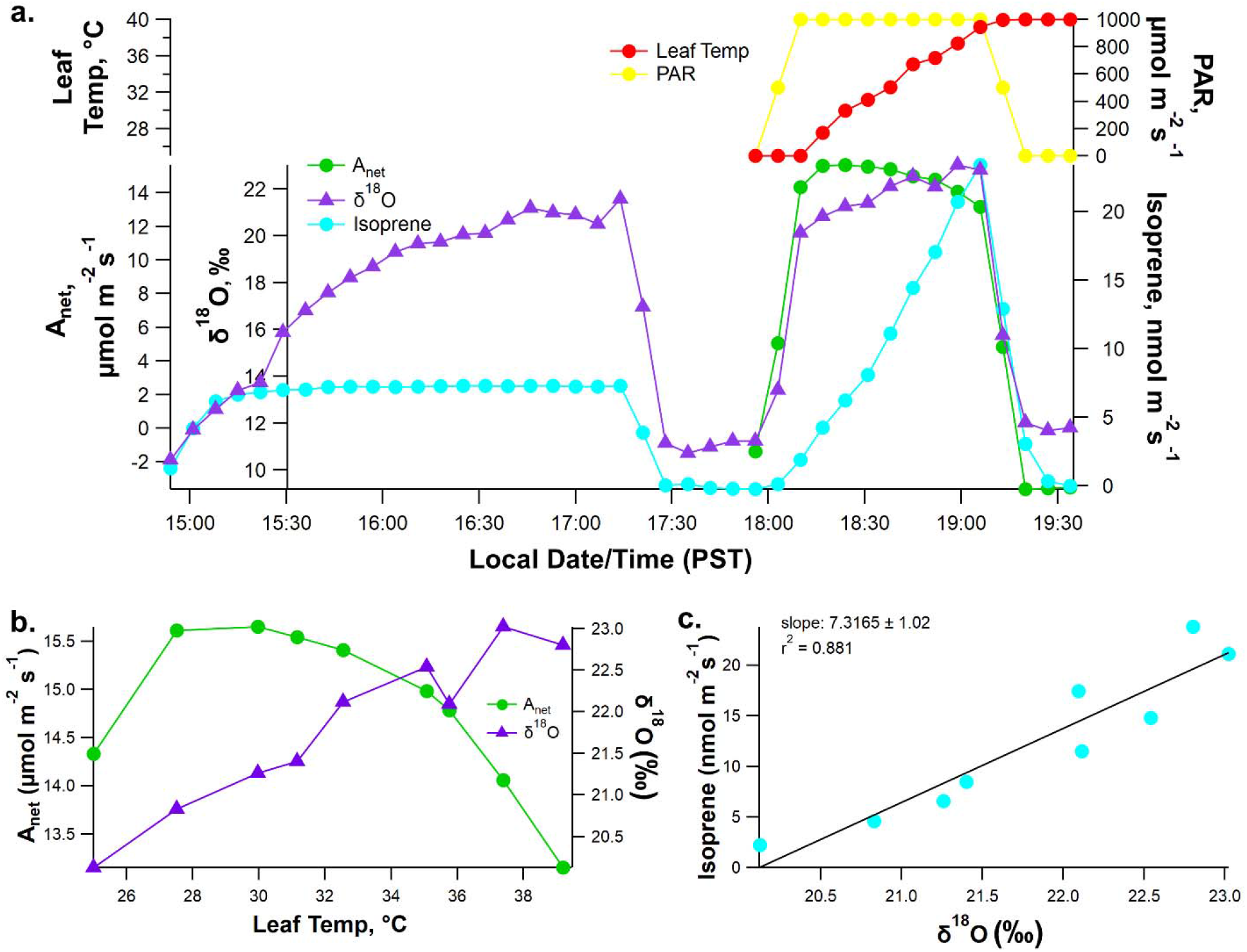
Example dynamics of ^18^O-labeled O_2_ evolution in the light as a function of leaf temperature from a detached poplar leaf pre-treated with ^18^O-water (**δ**^18^O 8,000 ‰) using the 36 cm^2^ leaf chamber. Pretreatment occurred under constant PAR (1,000 µmol m^-2^s^-1^), leaf temperature (32 °C), and leaf enclosure headspace CO_2_ (400 ppm). **a**. Example, real-time leaf-gas exchange fluxes of A_net_ and isoprene emissions together **δ**^18^O in headspace O_2_ during a controlled leaf temperature response curves under constant leaf headspace enclosure CO_2_ (400 ppm). Following pre-treatment, the light was switched off (PAR 0 µmol m^-2^s^-1^) and the leaf temperature reduced to 25 °C. Following measurements of dark gas parameters, PAR was returned to 1,000 µmol m^-2^s^-1^ and the leaf temperature response curve was initiated (25-40 °C). Finally, the light was switched off to determine the dark gas parameters at 40 °C leaf temperature, **b**. A_net_ and **δ**^18^O of O_2_ plotted as a function of leaf temperature, **c**. Linear regression between isoprene emissions and **δ**^18^O of O_2_ across leaf temperature in the light (PAR, 1,000 µmol m^-2^s^-1^). Note the slope of the regression as well as the 1:1 line shown.

### Optimal temperature of photosynthetic parameters

Data on optimal temperature (T_opt_) of A_net_, NOP, GOP, ETR, and isoprene emissions were compiled from the controlled temperature response curves using the small (N = 8) and large (N = 7) leaf chambers as well as the large leaf chamber during ^18^O-water labeling (N=7). As summarized in **Table 2**, A_net_ and NOP showed mean optimal temperatures of 31.0 +/- 3.1 °C and 31.0 +/- 3.4 °C, respectively. ETR and GOP showed a distinctively higher temperature optima of 35.0 +/- 1.8 °C and 34.9 +/- 1.8 °C, respectively. Isoprene emission had the highest temperature optimum at 38.9 +/- 1.0 °C. Despite a suppression in stomatal conductance at high temperature (g_s_ temperature optima of 33.0 +/- 5.7 °C) **(Figure 7**), transpiration (T_opt_ of 38.9 +/- 2.6) continued to increase at high leaf temperature.

**Figure 7.**
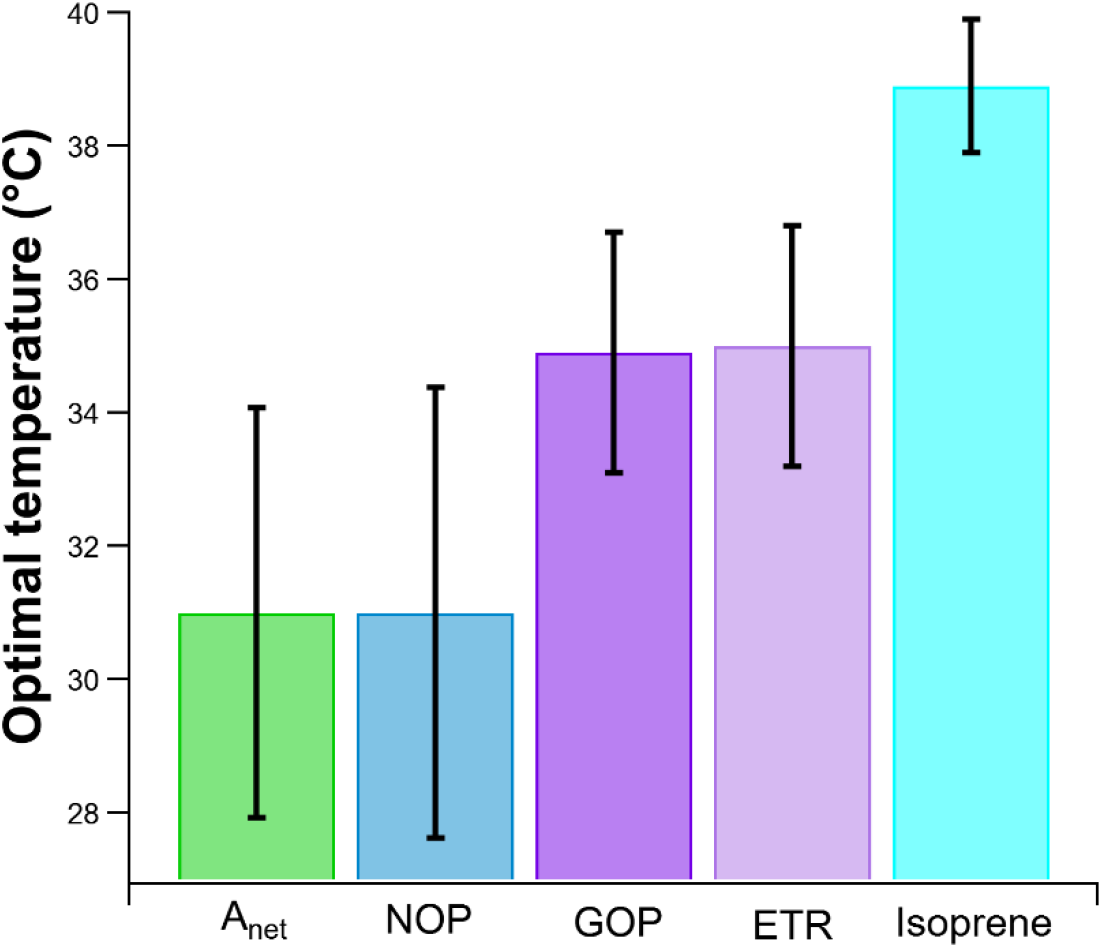
Average optimal temperature of net CO_2_ assimilation (A_net_), net oxygen production (NOP), gross oxygen production (*GOP), photosynthetic electron transport rates (**ETR), and isoprene emissions during controlled leaf temperature response curves (N = 15) using the small (N = 8) and large (N = 7) leaf chambers. Vertical error bars represent +/- 1 standard deviation. *GOP was only determined in the large chamber **ETR was only determined in the small chamber

**Table 2:** Optimal temperature (T_opt_) of leaf gas exchange and chlorophyll fluorescence parameters determined from the leaf temperature response curves.

| Parameter | T<sub>opt</sub> (°C) | Symbol (unit) | Instrument |
| --- | --- | --- | --- |
| Net photosynthesis | 31.0 ± 3.1 | A<sub>net</sub> (μmol m<sup>-2</sup> s<sup>-1</sup>) | Li6800 |
| Net oxygen production | 31.0 ± 3.4 | NOP (μmol m<sup>-2</sup> s<sup>-1</sup>) | CRDS in O<sub>2</sub> conc. mode |
| Stomatal conductance | 33.0 ± 5.7 | g<sub>s</sub> (mol m<sup>-2</sup> s<sup>-1</sup>) | Li6800 |
| Photosynthetic Electron Transport Rate | 35.0 ± 1.8 | ETR (μmol e<sup>-</sup> m<sup>-2</sup> s<sup>-1</sup>) | Li6800 |
| Gross oxygen production | 34.9 ± 1.8 | δ<sup>18</sup>O in O<sub>2</sub> | CRDS in δ<sup>18</sup>O mode |
| Transpiration | 38.9 ± 2.6 | E (mmol m<sup>-2</sup> s<sup>-1</sup>) | Li6800 |
| Leaf isoprene emissions | 38.9 ± 1.0 | Isoprene (nmol m<sup>-2</sup> s<sup>-1</sup>) | PTR-MS |

## Discussion

### Pros and cons of coupling gas exchange to O_2_ and isoprene flux measurements

In this study, we integrated real-time gas exchange fluxes of O_2_ (CRDS) and isoprene (PTR-MS) with a commercial photosynthesis system for leaf H_2_O and CO_2_ fluxes and chlorophyll fluorescence under controlled environmental conditions (Li-6800 with the 6 cm^2^ leaf fluorimeter chamber). This configuration provides a more complete picture of reactions of photosynthesis in leaves by simultaneously quantifying net oxygen production (NOP), net CO_2_ assimilation (A_net_), the assimilatory quotient AQ (A_net_/NOP), electron transport rate (ETR), and leaf isoprene emissions, which are dependent C_3_ carbon skeletons from the Calvin cycle and ATP and NADPH from ‘light’ reactions (Rodrigues et al., 2020). We optimized the air flow rate through the small leaf chamber (6 cm^2^) to enhance the O_2_ concentration gradient between reference and leaf chamber air while ensuring sufficient excess flow for CO_2_/H_2_O analysis (sample channel) by the IRGA as well as O_2_ and isoprene measurements by CRDS and PTR-MS, respectively. This allowed us to measure NOP concurrently with CO_2_, H_2_O, and isoprene fluxes (as well as other parameters such as ETR) (**Figure 1a**).

However, the drawback of this configuration is that small leaks that are unavoidable (between the leaf and the chamber gaskets) reduce excess air flow for IRGA analysis. Since external CRDS and PTR-MS sensors for O_2_ and isoprene emission measurements require a fixed flow rate, excessive leaks could decrease the flow below the minimum required for IRGA analysis (e.g. minimum air flow of 100-150 mL min^-1^ needed for CO_2_ and H_2_O mole fraction determination). In other words, a disadvantage of the small chamber is the need to carefully monitor flow rates after the leaf has been placed in the chamber, to verify that sufficient air flow exiting the leaf chamber arrives at the IRGA. In contrast, in the large chamber, there was a higher air flow rate and therefore, sufficient air flow from the leaf chamber to the IRGA was always achievable. Since the large leaf chamber had a larger volume than the small chamber (193.7 mL versus 87.3 mL), the residence time of circulating air increased from 0.26 min in the small chamber to 0.36 min in the large chamber (despite the increase in air flow through the large chamber). This allowed for the generation of a larger drawdown in O_2_ mole fraction between reference and outlet air. ΔO_2_ values reached up to 40 +/- 6 ppm using the small leaf chamber whereas ΔO_2_ values reached up to 139 +/- 52 ppm using the large leaf chamber. This feature makes the large chamber ideal for studying environmental dependencies of leaf O_2_ fluxes and ^18^O-labeling of O_2_ during photosynthesis under ^18^O-water. The main disadvantage of the large leaf chamber is the lack of an integrated fluorimeter needed to make measurements of chlorophyll fluorescence simultaneously with leaf gas exchange observations. Thus, we conducted replicated leaf response curves to light, CO_2_, and temperature using small and large leaf chambers. In contrast, we used the large leaf chamber exclusively with ^18^O-water labelling studies so as to have a better precision for δ^18^O measurements.

### Apparent assimilatory quotient (AQ)

Across light, CO_2_, and temperature response curves, A_net_ and NOP were tightly coupled and highly positively correlated, enabling the determination of apparent AQ values (AQ = A_net_/NOP) for each leaf experiment. For all environmental response curves, A_net_ displayed a slightly higher magnitude compared with NOP during conditions of low or negative net photosynthesis rates (dark, low C_i_, low temperature) and slightly higher magnitude during conditions of high net photosynthesis rates (e.g. saturating light and CO_2_ and optimal leaf temperature) (**Figs. 2-4**). This caused the values of AQ to be slightly higher than 1.0. When all leaf response curves were analyzed for AQ values (**Table 1**), no statistically significant difference was observed between AQ values determined from the light and C_i_ (two-tailed *P* value of 1.0000) or temperature and C_i_ (two-tailed *P* value of 0.4097) response curves. This suggests that the apparent leaf assimilatory quotient AQ does not depend on instantaneous changes in light, C_i_ (driven by changes in leaf headspace CO_2_ concentrations), or leaf temperature.

A recent review paper concluded that due to the lack of data on leaf O_2_ fluxes, the apparent assimilatory quotient (AQ) in the light is poorly characterized (Tcherkez et al., 2017). Leaf AQ values close to 1.0 suggest that O_2_ production during photosynthesis is balanced by CO_2_ assimilation and that photorespiratory and respiratory O_2_ consumption is balanced by CO_2_ production (Holloway-Phillips, 2018). However, AQ may deviate from 1.0 when significant activity of alternative electron transport processes occurs, not involved in CO_2_ fixation including nitrate photo-assimilation (Bloom et al., 1989) and lipid biosynthesis (Stumpf et al., 1963). For example, when wheat seedlings were grown with NH_4_^+^, leaf AQ values were 1.21 +/- 0.06. In contrast, seedlings grown with NO_3_^-^ showed significantly reduced AQ values of 1.13 +/- 0.05 (Smart and Bloom, 2001). Given that the availability of NH_4_^+^ as a nitrogen source in both the soil and daily watering in the current study, a significant reduction in AQ due to nitrate photo-assimilation in poplar leaves in this study is not expected. Thus, the AQ values determined from poplar leaves (1.27 +/- 0.12) compare well with AQ values (1.21 +/- 0.06) determined for wheat leaves supplied NH_4_^+^ (Smart and Bloom, 2001). Additional studies with wheat and maize using NH_4_^+^ as the nitrogen source also observed similar leaf AQ values in the light (e.g. 1.0-1.3), but observed some values less than 1.0 (e.g. 0.8) (Cousins and Bloom, 2004). Although these studies focused on the change in AQ between NO_3_^−^ versus NH_4_^+^ as a nitrogen source (ΔAQ) as a non-invasive measure of shoot NO_3_^−^ assimilation, a suppression of AQ as a function of light and leaf C_i_ was generally observed. This stands in contrast to our light (**Figure 3** and supplementary **Figures S1-S10**) and C_i_ response curves (**Figure 4** and supplementary **Figures S11-S24**) which showed a highly linear correlation between A_net_ and NOP, despite large changes in PAR and reference CO_2_. Similarly, no obvious suppression of AQ was observed as a function of temperature, which also showed highly linear correlations between A_net_ and NOP as leaf temperatures varied from 25-40 °C (**Figure 5** and supplementary **Figures S25-S54**). One potential explanation for the difference may be related to the method of determining AQ. For example in Cousins and Bloom, 2004, whom also used a dynamic leaf chamber, AQ was calculated by dividing the absolute flux of A_net_ and NOP at each light and C_i_ level, independent of the rest of the data in the response curve. Here, we determined the AQ value statistically by determining the slope of the regression between A_net_ and NOP across the entire PAR, C_i_, and leaf temperature response curves.

Large changes in light, C_i_, and leaf temperature can dramatically influence major metabolic processes other than photosynthesis that are involved in leaf CO_2_ and O_2_ metabolism including mitochondrial respiration (Souza et al., 2021) and photorespiration (Voss et al., 2013). However, because CO_2_ production is balanced by O_2_ consumption in the overall reactions, no net change in AQ is anticipated due to changes in their activities. Moreover, significant consumption of photosynthetic O_2_ by mitochondrial respiration in non-photosynthetic leaf cells and leaf, petiole, and stem vascular tissues could theoretically reduce NOP values determined from leaf gas exchange measurements by acting as a sink for photosynthetic O_2._ Leaf carbohydrate respiration by non-photosynthetic cells like epidermal cells, which make a small but significant (2%) component of total leaf respiration (Long et al., 2015), and petiole conduction bundles are often characterized by high rates of O_2_ uptake (Shugaeva et al., 2007). However, the delivery of CO_2_ to leaves via the transpiration stream from respiration in vascular tissues is expected to also reduce A_net_ as determined from leaf gas exchange. Thus, while A_net_ and NOP may be impacted by the delivery of respiratory CO_2_ to leaves via the transpiration stream and consumption of photosynthetically produced O_2_ by vascular tissues, AQ is not anticipated to deviate from 1.0 due to these processes. Moreover, photoreduction of O_2_ in photosynthetic leaf cells in the light yielding H_2_O_2_ that is rapidly converted back O_2_ and H_2_O can also be considered (Asada, 1999) (Asada, 1999). Given that this so called “water-water cycle” involves no net production or consumption of O_2_, changes in its activity are not expected to influence the assimilatory quotient AQ. Finally, despite the possibility that photosynthetic electron transport associated with chloroplastic lipid synthesis (e.g. isoprene) may alter AQ values (Stumpf et al., 1963), we found no evidence that conditions promoting high isoprene synthesis, suppresses AQ as has been observed with nitrate-photo-assimilation (Bloom et al., 1989). High rates of isoprene synthesis and emission at high light (**Figure 3** and supplementary **Figures S1-S10**) and temperature (**Figure 5** and supplementary **Figures S25-S54**), did not appear to alter the linear relationships between A_net_ and NOP. In contrast, the observations show a highly linear nature of A_net_ vs NOP for all light and temperature curves in this study, suggesting that isoprenoid synthesis represents a minor fraction of total ETR, and does not significantly suppress AQ values. This is consistent with a previous study that found that high temperature-stimulated isoprene emissions in a tropical broadleaf species represented only a small fraction (0.04%) of ETR (Rodrigues et al., 2020).

AQ values higher than 1.0 are difficult to explain, as they suggest that net CO_2_ assimilation is slightly higher than net O_2_ production. As first presented in studies focusing on the impact of nitrate photo-assimilation of leaf AQ values (Bloom et al., 1989; Cousins and Bloom, 2004), few studies have reported AQ values using dynamic plant chambers that allow for continuous quantification of net CO_2_ and O_2_ fluxes together under leaf headspace atmospheres maintained near natural ambient conditions, or during environmental response curves (Scafaro et al., 2017). The static method for plant-atmosphere gas exchange studies has limited temporal resolution, suffers from water condensation issues, requires a leak-free enclosure, which is often difficult to verify, rapidly generates unnatural atmospheres with excessive CO_2_ accumulation and O_2_ depletion, and requires physically removing the chamber or flushing it with fresh air before starting each measurement (Jardine et al., 2023). Nonetheless, AQ values of plant tissues based on gas-exchange methods are well known to be difficult to accurately obtain with a high degree of confidence because of the separate analytical techniques required (Scafaro et al., 2017). Systematic errors in either CO_2_ flux or O_2_ flux measurements will propagate into AQ values. In studies of respiration quotients (RQ) in the dark, when different methods to measure CO_2_ and O_2_ fluxes were compared, statistically significant differences were found between them. For example, when three methods to determine leaf dark respiration by fluorophore, O_2_-electrode, IRGA, and membrane inlet mass spectrometry techniques were compared, substantially different values were obtained (Scafaro et al., 2017). Using leaf dark respiration based on IRGA observations of net CO_2_ fluxes in the dark, RQ values equal to 1.0 as well as substantially less and greater than 1.0 could be obtained, depending on the method used for net O_2_ fluxes.

Rigorous calibration of CO_2_ and/or O_2_ sensors can improve the accuracy of flux measurements, but require a suite of highly accurate and precise gas standards. In our study, both sensors were factory calibrated within less than one year of the measurements. However, a full multipoint calibration of both CO_2_ and O_2_ sensors is necessary to either validate the factory calibrations or demonstrate a small offset between factory versus actual calibrations of the sensors. CO_2_ standards with high accuracy and precision are routinely prepared by specialized laboratories (e.g. 400 +/- 0.1 ppm). During the factory calibration of the Li6800 photosynthesis system used in this study, a set of CO_2_ standards spanning the range of concentrations encountered in the dynamic leaf chamber was conducted by the manufacturer (Licor Inc, Nebraska, USA). In contrast, to our knowledge, a high precision 21% O_2_ standard with precision of +/- 1 ppm or better is not available. Our method for AQ determination is highly dependent on an accurate CRDS calibration of O_2_ concentrations, with a particular importance on the slope of the instrument response to small changes (e.g. 0-200 ppm) in O_2_ concentrations (the sensitivity). A slight underestimate of the actual CRDS sensitivity to changes in O_2_ concentrations relative to the factory calibration would lead to underestimating NOP, and therefore overestimating AQ. Future studies should therefore attempt to address this issue by regularly calibrating both CO_2_ and O_2_ sensors with a range of high accuracy and precision gas standards that span the range of concentrations encountered in dynamic plant enclosures.

### Pros and cons of the ^18^O-labeling method

One of the grand challenges in terrestrial ecosystem science is daytime partitioning of net fluxes of CO_2_ and O_2_ into gross fluxes, i.e., (photo)respiration and gross photosynthesis (Wohlfahrt and Gu, 2015; Tramontana et al., 2020). At the leaf level in the light, using the Loreto method, gross photosynthesis under a ^13^CO_2_ atmosphere is directly quantified by the uptake flux of ^13^CO_2_ in parallel with gross (photo)respiration flux separately determined from the emission of ^12^CO_2_ (Pinelli and Loreto, 2003). More recently, a method enabling the partition of NOP fluxes into gross oxygen production (GOP) and (photo)respiratory O_2_ consumption has been used (Gauthier et al., 2018). In that study, the δ^18^O value of O_2_ in air of a detached leaf pre-treated with H_2_^18^O was used to estimate GOP, while NOP was determined separately using the O_2_/N_2_ ratio. Also, this method has been described as being highly technical (Holloway-Phillips, 2018), requiring an isotope ratio mass spectrometer (IRMS) to measure δ^18^O and O_2_/N_2_ (to estimate O_2_ mole fraction), a CRDS for water vapor concentration and δ^18^O of water vapor, and a separate CRDS for CO_2_ mole fraction. Nonetheless, this method was applied successfully during a controlled light response curve to partition NOP into GOP and (photo)respiration fluxes using French bean (*Phaseolus vulgaris* L.) leaves under 21% and 2% O_2_, but has not yet been applied to look at responses to leaf temperature, a key physiological variable for leaf net photosynthesis (Dusenge et al., 2019). Also, while a leaf chamber was used, a custom-made chamber was required to encapsulate both the leaf and the ^18^O-water container (thereby avoiding leaks). This custom configuration limited the control of environmental conditions that usual gas exchange systems are capable of, including air flow rate and humidity, light intensity and spectrum, leaf temperature and real-time, integrated CO_2_ and H_2_O mole fraction measurements (with IRGA) together with stomatal conductance to water vapor (g_s_).

Recent advances in high precision cavity ringdown spectroscopy (CRDS) of O_2_ have opened the door to terrestrial ecosystem O_2_ flux studies by directly resolving small changes in O_2_ concentrations in a background of ambient O_2_ concentrations (21%). For example, a O_2_ CRDS system developed by Picarro Inc. (G2207-i) has a precision of < 2 ppm O_2_ at 21% O_2_ with a 5-min average, enabling direct quantification of NOP fluxes from leaves. By switching the O_2_ CRDS into δ^18^O mode and feeding detached leaves with H_2_^18^O via the transpiration stream, we studied the temperature sensitivity of GOP using a method modified from Gauthier et al. (2018) allowing continuous gas exchange measurements of CO_2_, H_2_O, and isoprene (**Figure 1**). That is, unlike the method of Gauthier et al. (2018), which measured CO_2_ and H_2_O mole fraction and δ^18^O via two CRDS, here we obtained CO_2_ and H_2_O mole fraction using the integrated IRGAs of the gas exchange system. Also, δ^18^O was measured directly by the CRDS in isotope mode rather than IRMS, and we used the same CRDS (in concentration mode) to measure NOP (**Figure 1**).

If gross rates of oxygen production by photosystem II drive photosynthetic ETR, a tight correlation should be observed between GOP and ETR as leaf temperatures vary in the light. Consistent with this prediction, ETR, determined using the fluorimeter chamber, and δ^18^O of O_2_ during H ^18^O leaf labeling as a proxy for GOP, both increased to a similar optimal temperature of 35 °C before declining at higher temperatures. This is distinctly higher than the optimal temperature of NOP and A_net_ (31 °C). This suggests that the suppression of A_net_ and NOP at high temperature is mainly due to stomatal closure (T_opt_ of 33 °C) and accelerated rates of gross (photo)respiratory CO_2_ production/O_2_ consumption, rather than a decline in rates of gross photosynthesis. This is consistent with a model where at the optimal temperature for A_net_ and NOP of 31 °C in the light, relatively low rates of photorespiration, respiration, isoprenoid biosynthesis, and CO_2_/O_2_ recycling occur. In contrast at the optimal temperature for GOP and ETR of 35 °C, a reduction in g_s_ leads to a decrease in gross atmospheric CO_2_ uptake, which is partially compensated for by increased re-assimilation of internal CO_2_ sources derived from increasing temperature-dependent rates of photorespiration and mitochondrial respiration (Voss et al., 2013). In this metabolic model (**Figure 8**), suppression of A_net_ and NOP at 35 °C versus 31 °C is not due to high temperature stress of photosynthesis, but rather an acceleration of photosynthesis and (photo)respiration and their interactions via CO_2_ and O_2_ recycling (Garcia et al., 2019). In contrast, temperatures higher than 35 °C negatively impacted GOP and ETR, while still further accelerating (photo)respiration. Due to the decrease in O_2_ source (GOP) and increased O_2_ sinks via (photo)respiration, this results in further declines in A_net_ and NOP up to the highest leaf temperatures studied (40 °C).

**Figure 8:**
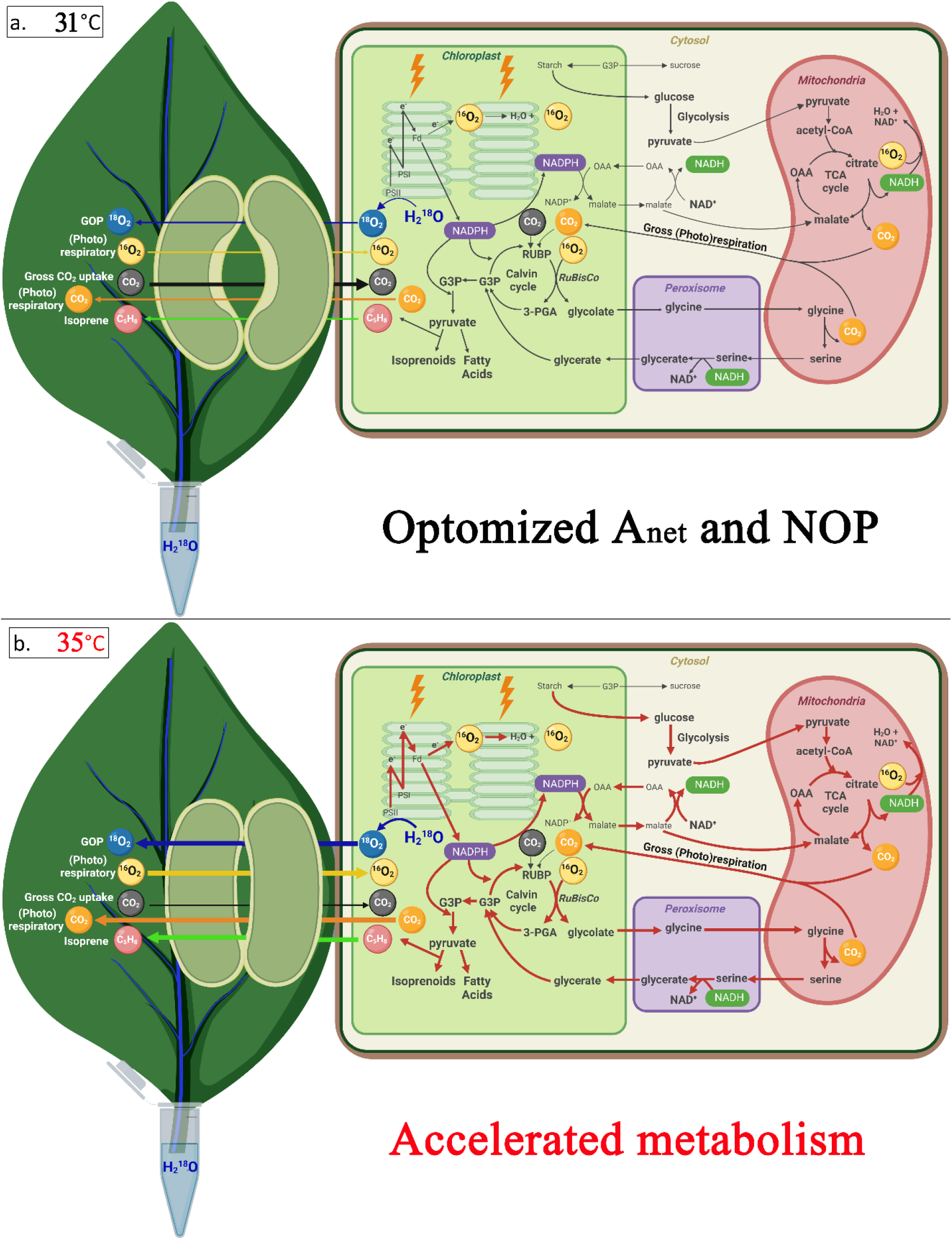
Simplified metabolic model of primary CO_2_ and O_2_ metabolism at 31 °C and 35 °C in poplar leaves. **a. 31.0 °C:** Optimal temperature for stomatal conductance (g_s_), A_nte_, and NOP in the light. Sub-optimal temperature for GOP and ETR. Low rates of photorespiration, respiration, isoprenoid biosynthesis, and CO_2_ re-assimilation. **b. 35 °C**: Suppression of g_s_ and gross atmospheric CO_2_ uptake. Optimal temperature for GOP and ETR. High rates of photorespiration, respiration, isoprenoid biosynthesis, and CO_2_ re-assimilation.

### Isoprene emission and its potential relationship with NOP and GOP

The response of leaf isoprene emissions to light (PAR), intercellular CO_2_ mole fraction (C_i_), and leaf temperature was broadly consistent with the common assumption of isoprene energetics models (also referred to as Niininets models) that isoprene emission relies on available reducing power in chloroplasts (Morfopoulos et al., 2013). This assumption reflects situations where the ATP and NADPH demand by the Calvin cycle for CO_2_ assimilation must be met before significant ATP/NADPH excess can be utilized for other pathways, such as the MEP pathway for isoprenoid biosynthesis (Rodrigues et al., 2020). Thus, one may anticipate that at elevated C_i_, a suppression of isoprene emissions together with a stimulation of A_net_ and NOP occurs, due to the increased demand for photosynthetic ATP/NADPH by the Calvin cycle (Morfopoulos et al., 2014), and this is what we observed (**Figure 4**). Similarly, at low light, isoprene emissions were barely detectable despite significant NOP and A_net_ fluxes, probably due to limited excess ATP/NADPH. Conversely, under light saturating conditions, further increases in PAR did not significantly enhance A_net_ or NOP, but strongly stimulated isoprene emissions, likely due to increased ATP/NADPH availability (**Figure 3**). Consequently, as predicted from isoprene photosynthesis energetic models and previous experimental observations (Jardine et al., 2016), the fraction of carbon (in % of A_net_) emitted as isoprene increased with light intensity.

We also observed a progressive increase in isoprene emission with temperature (**Figure 5**), consistent with previous studies where isoprene emissions increased up to 40 °C in many species (Harley et al., 1999). This effect could be attributed to both thermal stimulation of ETR and temperature-dependent activity of isoprene synthase (Monson et al., 1992; Morfopoulos et al., 2013). As observed here in poplar, ETR is frequently reported to have a higher leaf temperature optimum than A_net_ (Sage and Kubien, 2007). Also, isoprene energetic models predict a temperature optimum of isoprene emission that is strongly influenced by both optimal temperature of ETR and optimal temperature for isoprene synthase activity, which has been reported to be as high as 45 °C (Monson et al., 1992). Therefore, at leaf temperatures higher than the ETR and GOP optimum (35 °C), the increase in isoprene emissions up to 40 °C could be explained by a high temperature optimum for isoprene synthase (e.g. 45 °C) (Monson et al., 1992; Morfopoulos et al., 2013). However, we note that C_i_ decreased at leaf temperatures higher than the optimal for stomatal conductance (i.e., 33.0 °C) due to partial stomatal closure (Supplementary **Figure S69**). We thus suggest that in addition to the effect of isoprene synthase thermal optimum, the increase in isoprene emission was also driven by lower ATP/NADPH utilization for carboxylation in the Calvin cycle, resulting from partial stomatal closure and the decline in C_i_.

Taken as a whole, our results are in excellent agreement with the availability of NADPH and ATP in the chloroplast being rate-limiting for isoprenoid biosynthesis (Rasulov et al., 2009; Rasulov et al., 2018). While carbon limitations for isoprene biosynthesis have been generally considered negligible, previous studies using CO_2_-free air demonstrated that isoprene emissions can occur at surprisingly high rates, are light and temperature stimulated, and associated with refixation of (photo)respiratory CO_2_ (Garcia et al., 2019). In Garcia et al. (2019), the comparison of isoprene emission under CO_2_-free atmosphere with 400 ppm-CO_2_ atmosphere concluded that CO_2_ refixation could account for the “alternate carbon source for isoprene not from uptake of atmospheric CO_2_”. While non-photosynthetic stored carbon sources for isoprene have been discussed in the literature (Jardine et al., 2014), our observations are consistent with the idea that carbon limitation for isoprene occur only at very low C_i_ (Lantz et al., 2019). In other words, regardless of whether carbon atoms come from refixation or reserve remobilization, our results suggest an important role of carbon recycling in leaves to sustain isoprene emission, not only under photorespiratory conditions or high light but also high temperature, and thus recycling is perhaps an important thermotolerance mechanism.

### Conclusions and perspectives

Our study shows that coupling O_2_ and isoprene exchange to traditional CO_2_/H_2_O gas exchange is possible, using CRDS-based oxygen and PTR-MS-based isoprene measurements. This configuration allows a more complete picture of the photosynthetic redox budget via photosynthetic production of O_2_, electron transport rate (ETR), and isoprene biosynthesis. This opens avenues for useful measurements during photosynthesis, such as the assimilatory quotient (AQ). Also, our findings may help resolve some confusion in the literature as to whether isoprene emissions may or may not be directly linked to net photosynthesis. In effect, we found that isoprene emission can be uncoupled from A_net_, e.g., at high temperature or low C_i_ (**Figs. 5 and 6, and Supplementary Figures S1-S54**), and thus it is unlikely that isoprene biosynthesis strictly depends on photosynthesis rate or carbon provision by photosynthates. In contrast, when isoprene emission observed during the temperature response curve with ^18^O-water was plotted against δ^18^O (**Fig. 6**), there was a linear correlation (with a R^2^ of 0.88; see also **Supplementary Figures S55-S68**). Therefore, our results suggest that (*i*) isoprene emission is related to electron generation by photolysis and thus probably by excess photosynthetic ATP/NADPH (not consumed by the Calvin cycle), and (*ii*) is carbon-limited only when gross photosynthesis declines considerably. We nevertheless recognize that dual isotopic labelling with ^13^CO_2_ and ^18^O-water would be useful to ascertain this and quantify precisely the link between ^13^C-isoprene appearance and ^18^O_2_ evolution. This will be addressed in another study.

### Supplemental data section

The following supplemental materials are available in the online version of this article. All raw and derived leaf gas exchange and chlorophyll fluorescence data presented in **Figures 3-6** and supplementary **Figures S1-S68** with this manuscript are available to download free of charge at Mendeley Data (data.mendeley.com) (**doi web link**) and the Environmental System Science Data Infrastructure for a Virtual Ecosystem (ESS-DIVE, ess-dive.lbl.gov) data repository (**doi web link**).

**Figure S1-S6:** Biological replicates #1-3 of real-time leaf-gas exchange fluxes of A_net_, NOP, E, and isoprene emissions together with g_s_ and ETR during controlled light response curves (PAR) using the 6 cm^2^ leaf chamber.

**Figure S7-S10:** Biological replicates #1-2 of real-time leaf-gas exchange fluxes of A_net_, NOP, E, and isoprene emissions together with g_s_ during controlled light response curves (PAR) using the 36 cm^2^ leaf chamber.

**Figure S11-S18:** Biological replicates #1-4 of real-time leaf-gas exchange fluxes of A_net_, NOP, E, and isoprene emissions together with g_s_ and ETR during a controlled C_i_ response curve using the 6 cm^2^ leaf chamber.

**Figure S19-S24:** Biological replicates #1-3 of real-time leaf-gas exchange fluxes of A_net_, NOP, E, and isoprene emissions together with g_s_ during a controlled C_i_ response curve using the 36 cm^2^ leaf chamber.

**Figure S25-S40:** Biological replicates #1-8 of real-time leaf-gas exchange fluxes of A_net_, NOP, E, and isoprene emissions together with g_s_ and ETR during a controlled leaf temperature response curve using the 6 cm^2^ leaf chamber.

**Figure S41-S54:** Biological replicates #1-7 of real-time leaf-gas exchange fluxes of A_net_, NOP, E, and isoprene emissions together with g_s_ during a controlled leaf temperature response curve using the 36 cm^2^ leaf chamber.

**Figure S55-S68:** Biological replicates #1-7 of real-time leaf-gas exchange fluxes of A_net_ and isoprene emissions together with **δ**^18^O of headspace O_2_ during a controlled leaf temperature response curves under constant leaf headspace enclosure CO_2_ (400 ppm) and PAR (1000 µmol m^-2^ s^-1^) from a detached poplar leaf pre-treated with ^18^O-water.

**Figure S69**: Example dependence of C_i_ on leaf temperature during a leaf temperature response curve.

## Acknowledgements

We kindly thank Bryan Taylor at Lawrence Berkeley National Laboratory for the technical support.

## Author contributions

K. Jardine acquired funding and developed the illustrations and together with S. Som designed and configured the leaf gas exchange system; S. Som collected and analyzed all data, developed all data figures, and together with L. Gallo, archived the public data set; A. Sunder assisted with supplies procurement and statistical analysis; L. Gallo developed the TOC icon; J. Demus and C. M. Wistrom maintained optimal growth conditions of poplar trees in the UC Berkeley Oxford Tract greenhouse and facilitated the integration of a real-time volatile metabolomics laboratory; All authors assisted in manuscript preparation and revising the work critically for important intellectual content. In addition, T. F. Dominges and G. Tcherkez assisted with critical scientific insights and detailed manuscript revisions.

